# Honey bee (*Apis mellifera*) larval pheromones regulate gene expression related to foraging task specialization

**DOI:** 10.1101/587113

**Authors:** R Ma, J Rangel, CM Grozinger

**Author notes:** Corresponding Author: Rong Ma.

## Abstract

**Background:** Foraging behavior in honey bees (*Apis mellifera*) is a complex phenotype which is regulated by physiological state and social signals. How these factors are integrated at the molecular level to modulate foraging behavior has not been well-characterized. The transition of worker bees from nursing to foraging behavior is mediated by large-scale changes in brain gene expression, which are influenced by pheromones produced by the queen and larvae. Larval pheromones can also stimulate foragers to leave the colony to collect pollen, but the mechanisms underpinning this rapid behavioral plasticity are unknown. Furthermore, the mechanisms through which foragers specialize on collecting nectar or pollen, and how larval pheromones impact these different behavioral states, remains to be determined. Here, we investigated the patterns of gene expression related to rapid behavioral plasticity and task allocation among honey bee foragers exposed to two larval pheromones, brood pheromone (BP) and (E)-beta-ocimene (EBO).

**Results:** We hypothesized that both pheromones would alter expression of genes in the brain related to foraging and would differentially impact expression of genes in the brains of pollen compared to nectar foragers. Combining data reduction, clustering, and network analysis methods, we found that foraging preference (nectar vs. pollen) and pheromone exposure are each associated with specific brain gene expression profiles. Furthermore, pheromone exposure has a strong transcriptional effect on genes that are preferentially expressed in nectar foragers. Representation factor analysis between our study and previous landmark honey bee transcriptome studies revealed significant overlaps for both pheromone communication and foraging task specialization.

**Conclusions:** Social signals (i.e. pheromones) may invoke foraging-related genes to upregulate pollen foraging at both long and short time scales. These results provide new insights into how social signals integrate with task specialization at the molecular level and highlights the important role that brain gene expression plays in behavioral plasticity across time scales.

## Background

One of the hallmarks of insect sociality is division of labor, whereby group members specialize on different tasks that are essential to group survival and reproduction [1, 2]. Understanding the proximate and ultimate mechanisms mediating social behavior, division of labor, and task specialization, is a major focus of behavioral sociobiology [3–9]. Several studies have clearly demonstrated that complex animal behaviors, including social interactions, are regulated by transcriptional, neural, and physiological networks [10–13]. Others have suggested that behavioral ontogeny is mediated by differential regulation of core, well-conserved transcriptional or physiological “toolkits” that regulate behavioral modules [4, 14–20]. However, the mechanisms mediating more rapid shifts in behavior and task specialization have not been examined as thoroughly [21–23].

Like many social insects, honey bee (*Apis mellifera*) workers exhibit a form of age-based task allocation in which their behavioral repertoires incrementally expand or shift over the course of an individual’s lifetime [24]. This phenomenon—called age-based polyethism—is regulated both genetically and environmentally, and provides a tractable system in which to investigate temporal dimensions of behavioral plasticity [25, 26]. Honey bees spend the first weeks of their lives performing tasks within the relative safety of the hive, including tending to the needs of developing larvae (i.e. nursing), before transitioning to increasingly dangerous tasks near the nest entrance and beyond, including foraging [27]. Once they begin foraging, workers may further specialize by collecting predominantly one floral resource type, either pollen or nectar [28], and their proclivity for pollen versus nectar foraging can persist throughout their lives. Bees that specialize on nectar versus pollen foraging exhibit distinct behavioral, physiological, and transcriptional traits. For example, upon returning to the colony, nectar foragers regurgitate collected nectar to nestmates waiting to process it, while pollen foragers pack their pollen loads into honeycomb themselves [29, 30]. Nectar and pollen foragers also differ in neural and sensory responses to sugar [31] and pheromones [32, 33].

Pheromone communication in honey bees plays a key role in behavioral transitions across time scales [10, 34–37]. Pheromones are typically categorized by the time scale at which they induce behavioral changes: primer pheromones cause slow, enduring changes in physiology in receivers, while releaser pheromones cause rapid, ephemeral responses in receivers. Primer pheromones generate these long-term changes in behavior and physiology by altering patterns in gene expression, especially in the brain [10, 34–37]. Brood pheromone and queen pheromones, for example, both delay the behavioral transition from nurses to foragers by altering the expression of large numbers of genes in worker brains [34, 37]. Releaser pheromones elicit rapid behavioral changes either by activating or modulating neural circuits, triggering molecular signaling pathways, or regulating gene expression [35, 38–41]. For example, alarm pheromone in honey bees elicits aggressive behaviors against intruders by activating the expression of immediate early genes in the brain [35], while one component of queen pheromone, homovanillyl alcohol, elicits grooming behavior from workers by binding to an olfactory receptor in antennae, activating dopamine receptors in the brain, and regulating brain gene expression [34, 42, 43].

Honey bee larval pheromones cause primer and releaser effects that blur the distinction between these categories, which provides a fascinating opportunity to understand regulation of behavior across time scales. Two larvae-produced pheromones, brood pheromone (BP) and (E)-beta-ocimene (EBO), have been shown to elicit rapid increases in foraging within an hour of exposure, presumably by stimulating foraging behavior in a colony’s existing foragers. The effect of brood pheromones on forager behavior seems to be driven by an increase in pollen foraging specifically [44]. Both pheromones also increase the foraging force of the colony, over the long-term, by accelerating the transition of bees from performing within-hive roles to foraging [45–47]. This is a fascinating system because both pheromones regulate the same type of behavior, but at different temporal scales. How these behavioral transitions across different temporal scales are related, or how their mechanisms interact, remains to be determined.

In this study, we evaluated the transcriptional mechanisms underlying rapid changes in honey bee foraging behavior, and juxtaposed these rapid changes, triggered by social signals, with more stable differences in gene expression associated with task specialization. Given that foragers have similar behavioral responses to BP and EBO [48], we hypothesized that these two pheromones regulate a common set of foraging genes in the brain (i.e. a foraging “toolkit”). Because BP and EBO have more pronounced effects on pollen foraging than nectar foraging [44, 46], we further hypothesized that larval pheromones affect gene expression in pollen foragers more strongly than in nectar foragers. We thus compared the effects of EBO and BP exposure on foragers previously found to specialize on nectar or pollen to test the following four predictions: 1) foragers specializing on pollen versus nectar foraging exhibit distinct patterns of gene expression, 2) BP and EBO stimulate the same transcriptional profiles in the brains of forager bees 3) both larval pheromones have more pronounced effects on gene expression in pollen foragers than nectar foragers, and 4) changes in the same behavior at different time scales (i.e., the transition to and/or stimulation of pollen foraging) utilize similar molecular mechanisms.

Combining differential gene expression, clustering, and network analyses, our study presents several lines of evidence that show that larval pheromones and foraging are associated with expression profiles of common suites of genes, and that these genes are related to metabolic and fatty acid biosynthesis pathways and integral components of the membrane, including sodium channels. Our study elucidates the molecular mechanisms underlying task allocation and highlights the important role that brain gene expression plays in behavioral plasticity across time scales. It also probes the interface between ephemeral and more consistent changes in behavior to gain insight into mechanisms that permit behavioral plasticity and complexity across time.

## Results

### Transcript quantification

The RNA samples collected in this study were extracted from mushroom bodies of pollen and nectar foragers exposed to one of three pheromone treatments: paraffin oil control, brood ester pheromone (BP), or E-beta-ocimene (EBO) (Fig. 1). The number of RNA-seq reads per sample ranged from 41-94 million, with an average of 65 million reads per sample. After quality filtering and adapter trimming, an average of 69% of the reads per sample were pseudoaligned to generate transcript abundance for each annotated transcript in the recently updated honey bee genome annotation (Amel_HAv3.1). Overall, 9,179 genes were represented in the data set, representing 74% of 12,332 annotated honey bee genes.

**Figure 1.**
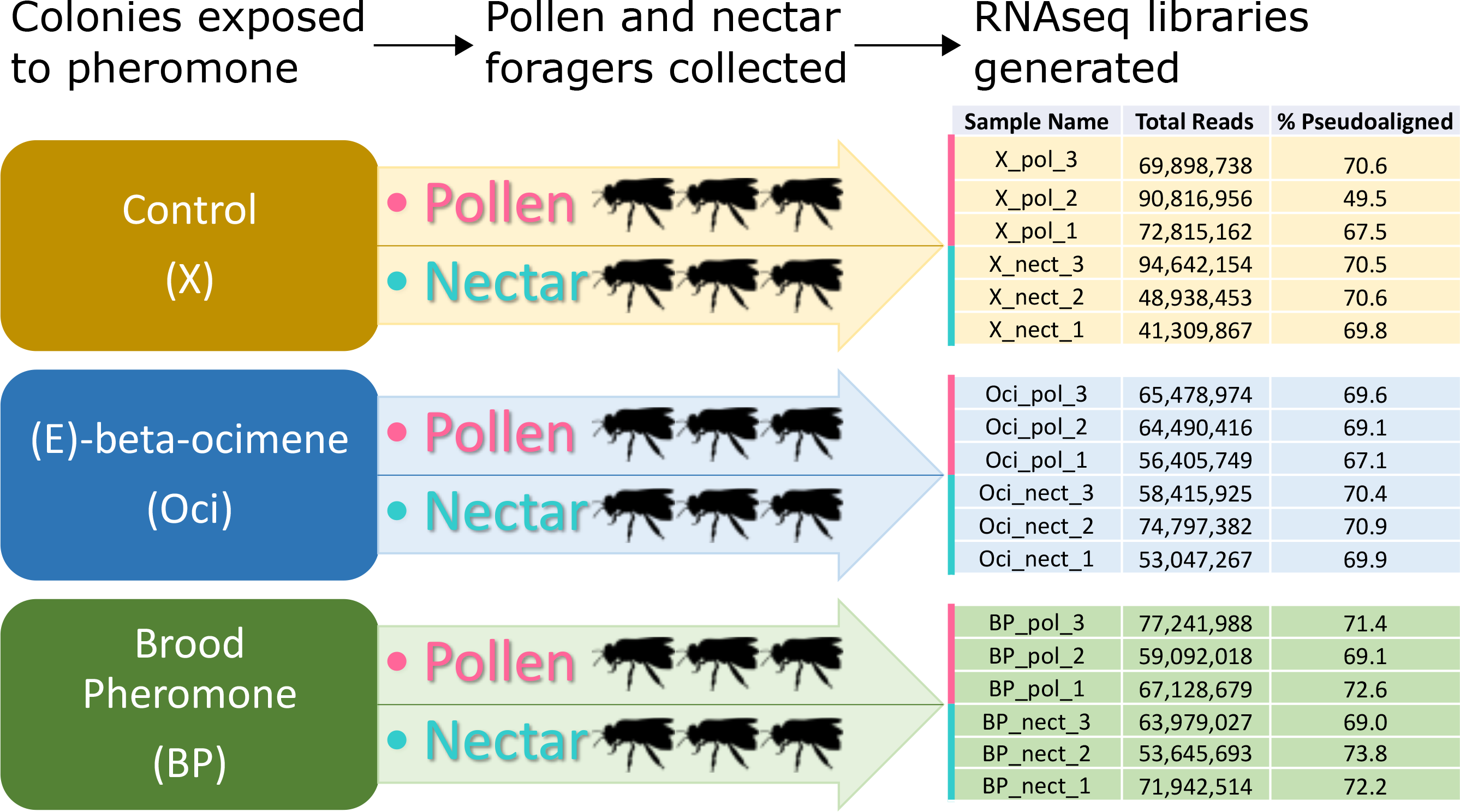
Overview of experimental design and sequencing. RNA-seq libraries were generated from nectar and pollen foragers exposed to three pheromone treatments. Three pooled pollen forager samples and three pooled nectar forager samples were collected for each pheromone treatment. Each bee diagram represents a sample, though two brains per used for each sample. Resulting numbers of reads per sample and percentages of those reads that mapped to honey bee genome are presented in a table to the right.

### Differential gene expression

Differential gene expression analysis was performed to characterize the effects pheromone treatment, forager-type, and the interaction between pheromone and forager type. There were 533 differentially expressed genes (DEG) whose expression varied in at least one contrast (FDR<0.05), including 269 DEG related to pheromone treatment and 326 DEG related to forager type (Table 1). Additionally, there were 131 DEG that showed a statistically significant interaction between forager type and pheromone treatment.

**Table 1:**
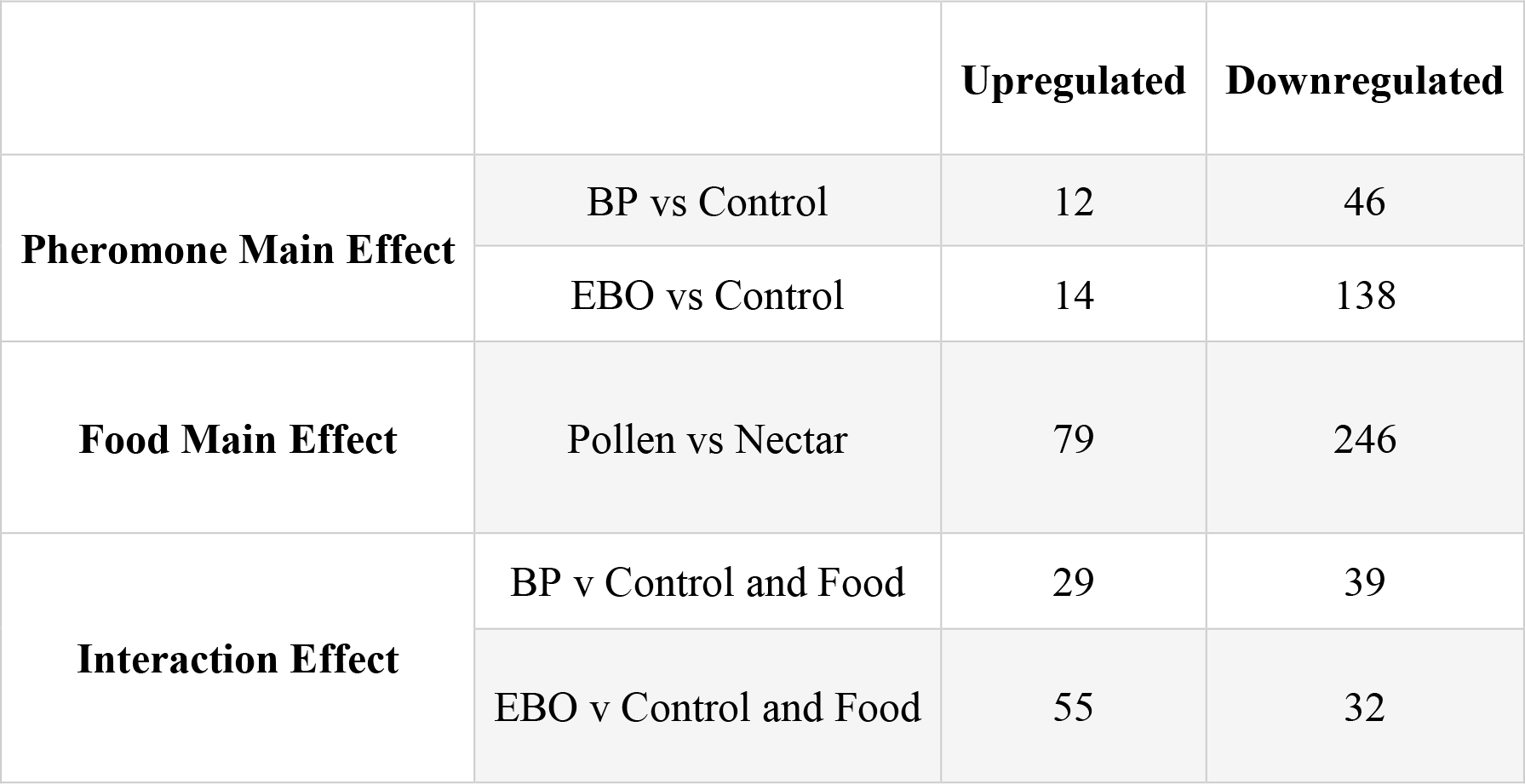
Numbers of DEG in all pairwise comparisons

|  |  | Upregulated | Downregulated |
| --- | --- | --- | --- |
| **Pheromone Main Effect** | BP vs Control | 12 | 46 |
|  | EBO vs Control | 14 | 138 |
| **Food Main Effect** | Pollen vs Nectar | 79 | 246 |
| **Interaction Effect** | BP v Control and Food | 29 | 39 |
|  | EBO v Control and Food | 55 | 32 |

Of the 269 DEGs related to pheromone treatment, there were 58 DEGs between BP and control samples, and 152 DEGs between EBO and control samples, indicating that EBO’s effect on gene expression was almost three times greater than that of BP. In addition, there were 148 genes that showed differences between BP and EBO samples. Because there were many genes that were differentially expressed in more than one contrast, we performed hypergeometric tests to determine whether there were more shared DEGs than those from random expectation among pheromone treatments, and between pheromone treatments and forager type. There were significant overlaps between all pairwise comparisons of pheromone treatment, indicating that BP and EBO regulate expression of a common subset of genes or genetic pathways (Table 2).

**Table 2.** Overlaps between pheromone-related DEG

| First Contrast | Second Contrast | DEGs in First Contrast | DEGs in Second Contrast | Overlap |
| --- | --- | --- | --- | --- |
| BP vs Control | EBO vs Control | 58 | 152 | 39* |
| *significantly greater overlap of genes than expected by chance; P<0.001; hypergeometric test |  |  |  |  |

Pheromone-related DEGs were then compared to DEGs that differed between nectar and pollen foragers, and there were significant overlaps between foraging-related and pheromone-related DEGs (Table 3). To further explore these results, we split the foraging-related DEGs into those that were upregulated in pollen foragers and those upregulated in nectar foragers, and again looked for overlaps with DEGs from each pheromone treatment. Interestingly, there were significant overlaps between pheromone-related DEGs and DEGs upregulated in nectar foragers (Table 4; hypergeometric tests, p<0.01), but not between pheromone-related DEGs and DEGs upregulated in pollen foragers. In summary, BP and EBO both regulated foraging-related genes, and this effect was driven primarily by genes upregulated in nectar foragers relative to pollen foragers.

**Table 3.** Overlaps between pheromone- and foraging-related DEG

|  | Pheromone genes | Food Genes | Overlap |
| --- | --- | --- | --- |
| BP vs Control | 58 | 386 | 41* |
| EBO vs Control | 152 | 386 | 71* |
| *significantly greater overlap of genes than expected by chance; P<0.001; hypergeometric test |  |  |  |

**Table 4:**
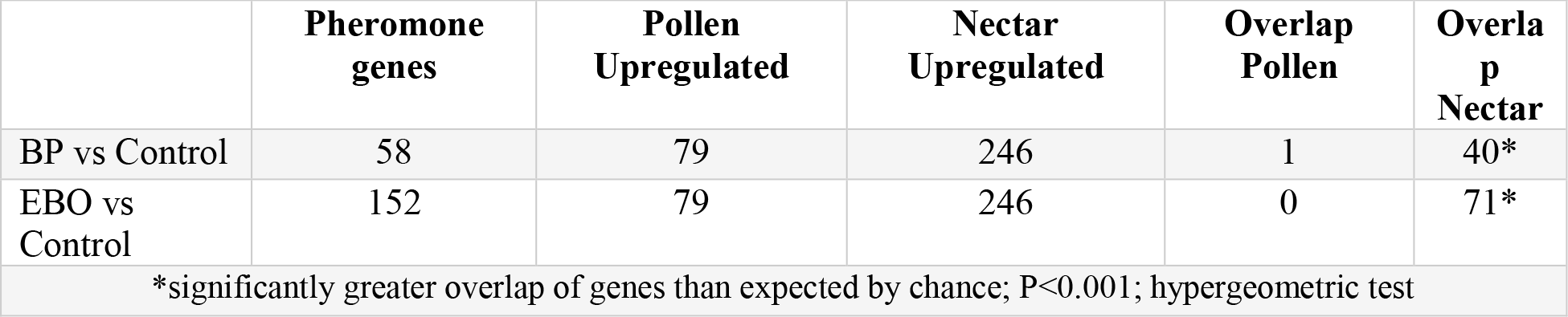
Overlaps between pheromone- and foraging-related genes

|  | Pheromone genes | Pollen Upregulated | Nectar Upregulated | Overlap Pollen | Overlap Nectar |
| --- | --- | --- | --- | --- | --- |
| BP vs Control | 58 | 79 | 246 | 1 | 40* |
| EBO vs Control | 152 | 79 | 246 | 0 | 71* |
| *significantly greater overlap of genes than expected by chance; P<0.001; hypergeometric test |  |  |  |  |  |

To better understand the function of differentially expressed genes associated with forager type and pheromone treatment, we performed gene ontology (GO) enrichment analysis for DEGs associated with pheromone treatment, forager type, and their interaction. DEGs associated with forager type were significantly enriched for GO terms related to lipid metabolism and trypsin-like serine proteases (FDR < 0.05). DEGs related to pheromone treatment were enriched for integral components of membrane, fatty acid metabolism, and lipid biosynthesis (FDR < 0.05). Finally, DEGs related to the interaction of pheromone treatment and forager type were enriched for lipid biosynthesis and metabolism (FDR <0.05).

The DEGs associated with either EBO or BP were also analyzed separately. Because there were few upregulated genes associated with either pheromone, up- and down-regulated genes for each pheromone were pooled during pathway enrichment analysis; however, it should be noted that the results for pheromone could potentially be driven by down-regulated genes. DEGs associated with BP exposure were enriched for lipid biosynthesis and integral components of the membrane (FDR < 0.05). DEGs associated to EBO exposure were enriched for integral components of membrane, fatty acid biosynthetic processes, fatty acid metabolism, and pentose phosphate pathway. There was a significant overlap of 39 genes between BP and EBO exposed foragers compared to controls (P<0.05), and these DEGs were significantly enriched for metabolic pathways and fatty acid metabolism (FDR<0.05).

### Hierarchical clustering and Principal Components Analysis (PCA)

Hierarchical clustering analysis and PCA were used to better understand broad patterns across all DEGs. Based on all variance-stabilized gene expression values of DEGs, hierarchical clustering grouped samples with identical combinations of pheromone treatment and forager type (Fig. 2) significantly more often than random expectation based on 10,000 iterations of multiscale bootstrap resampling (P<0.05; Supplementary Fig. 1). Nectar foragers exposed to either BP or control pheromone treatments clustered together; however, nectar foragers exposed to EBO clustered with pollen foragers, suggesting that EBO exposure resulted in gene expression patterns of nectar foragers that were more similar to those of pollen foragers. This is consistent with the observation that EBO had a greater effect on gene expression than BP. Pollen foragers exposed to BP or EBO were more similar to each other than either group was to pollen foragers exposed to control treatments. Genes were also clustered based on the similarity of their expression, and several large clusters of genes emerged.

**Figure 2.**
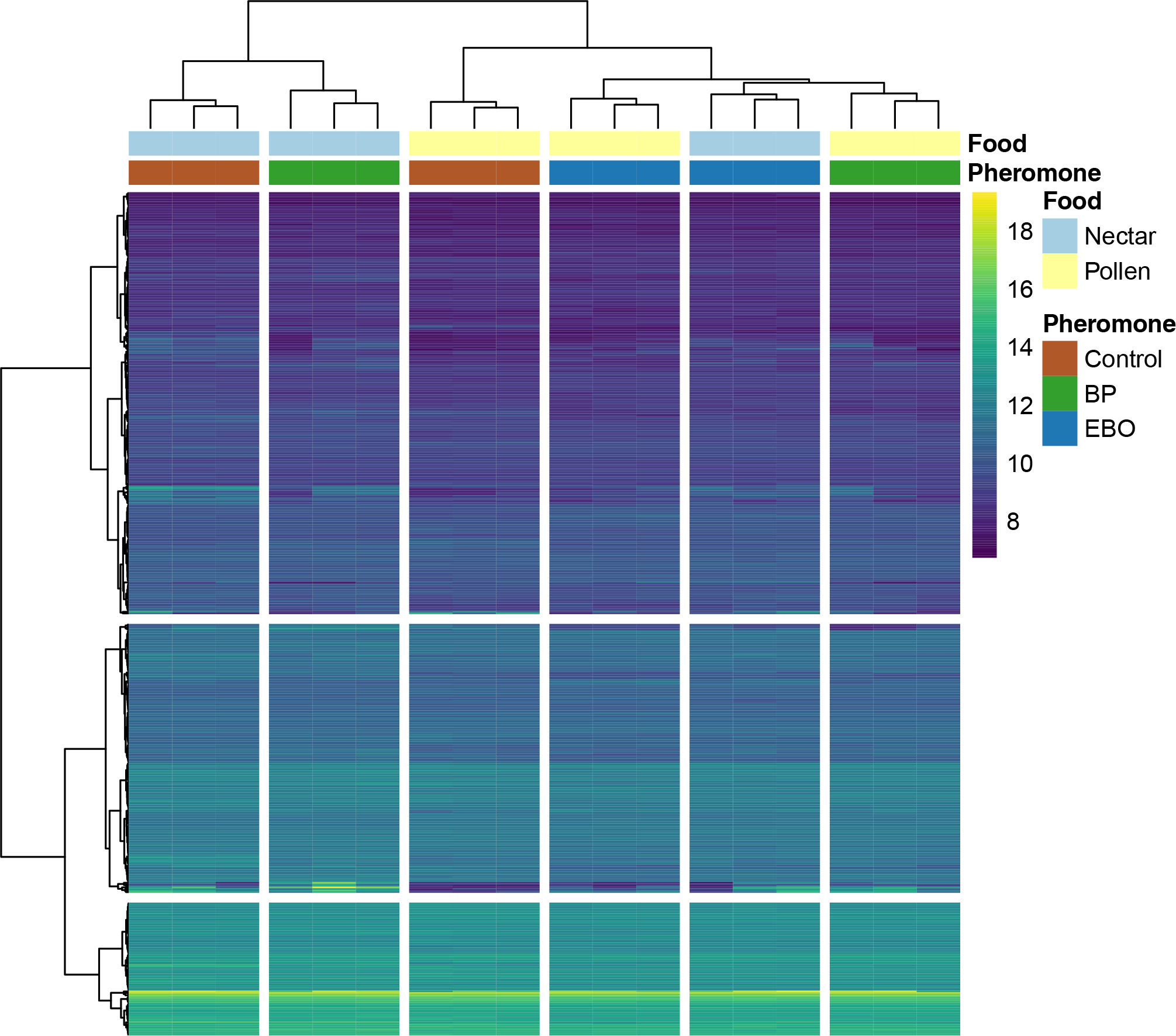
Heatmap for the hierarchical clustering of brain gene profiles. Honey bees foraging on pollen or nectar were exposed to pheromone treatments: Brood pheromone (BP), E-beta-ocimene (EBO), or a control. Rows correspond to differentially expressed genes, and columns represent samples. Food and pheromone treatments for each sample are represented between sample dendrogram and heatmap. The scale bar indicates z-scores of variance stabilized gene expression values, with highly expressed genes in lighter colors and lower expression in darker colors. Clustering of samples shows two branches main branches, which correspond broadly to nectar foraging (left) and pollen foraging(right); however, nectar foragers exposed to EBO have expression profiles more similar to pollen foragers. Within pollen and nectar branches, there is also a split in pheromone treatments.

To better understand the contributions of pheromone treatment and forager type on patterns of gene expression, we performed PCA on all differentially expressed genes with samples grouped by treatment. Each principal component (PC) was composed of a linear combination of many genes. Together, the first two PCs explained 63% of variance in the data, and the PCs were useful in separating samples by both pheromone treatment and forager type (Fig. 3). The first PC explained 46 % of variance and separates nectar and pollen foragers, indicating that the greatest axis of variation in gene regulation related to forager type. This is consistent with results from the differential gene expression analysis, which showed that there were more DEGs associated with forager type than with pheromone exposure. Nectar foragers generally exhibited more negative values in the PC1, while pollen foragers exhibited more positive values. The second principal component explained 17.0 % of the variance in the DEGs and began to separate pheromone treatment from each other, although the separation was less distinct than for forager type. Specifically, PC2 seemed to separate bees exposed to control pheromone treatment from those exposed to BP, while samples from bees exposed to EBO were more intermediate. Pollen foragers, especially those exposed to EBO and control treatments, seemed to have a lower variance than nectar foragers in both principal components.

**Figure 3.**
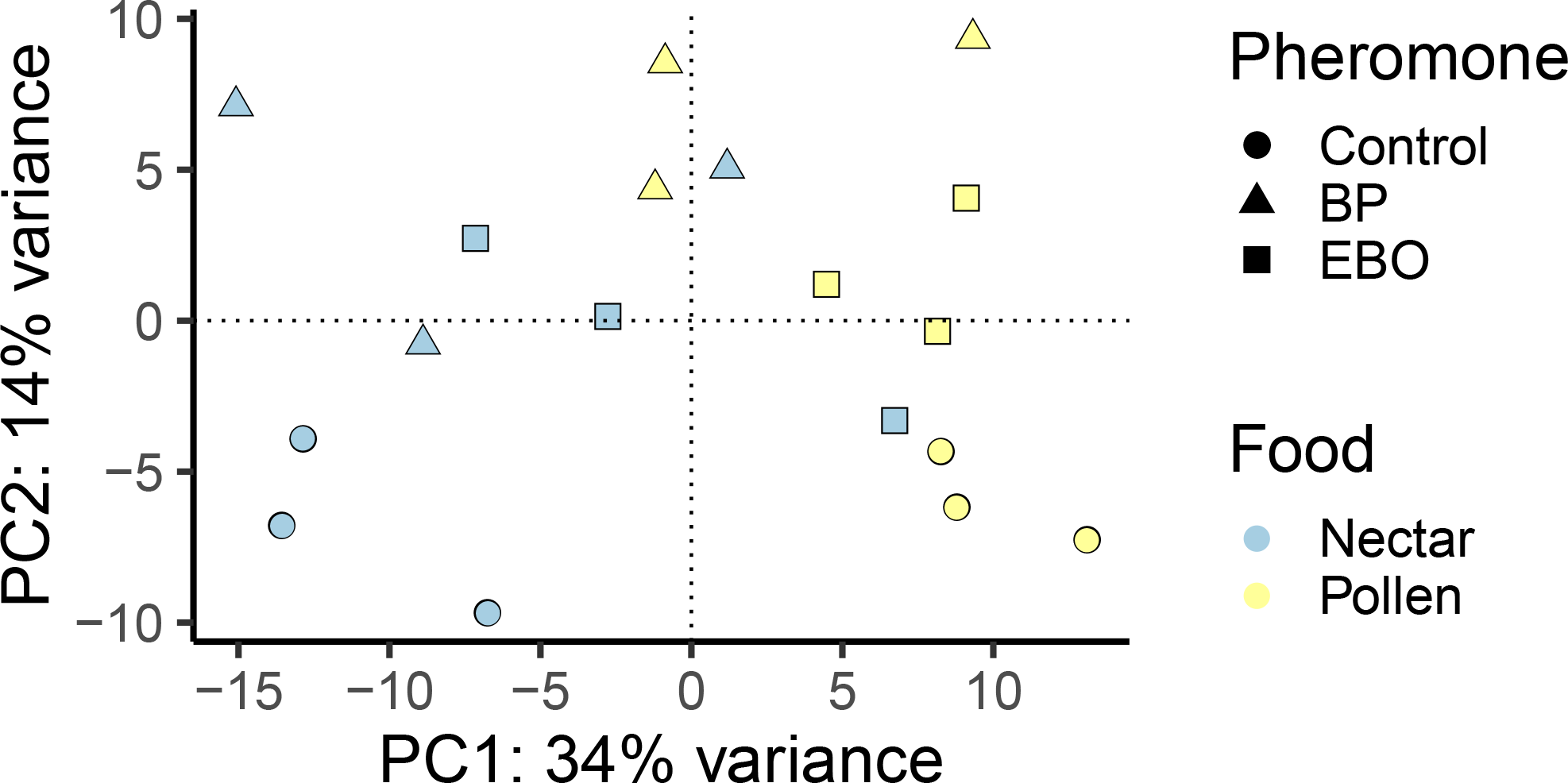
Principal component analysis of all DEG. The first two principal components (PCs) are displayed, together representing 63% of the total variation. Each point represents a single sample. PC1 separates samples based on food preference, whereas PC2 pheromone treatment, particularly for nectar foragers. Shape represents pheromone treatment. Color represents pollen or nectar forager-type. The percentage of variation in transcript expression patterns explained by each PC is shown in the y-axis.

### Overlaps with landmark studies

To explore the relationship between the results shown above and those of previous similar studies, we performed representation factor analysis between our results and landmark honey bee transcriptome studies (Tables 5, 6) [37, 49]. Whitfield et al. [49] identified DEGs related to foraging ontogeny, while Alaux et al. [37]. identified DEGs related to long-term exposure to BP (i.e. primer pheromone effects). We found a significant overlap between the foraging-related genes identified in our study and those identified by [49] (hypergeometric test, P<0.05;Table 6). Thus, genes that were differentially expressed in the brains of nectar and pollen foragers (our study) overlapped significantly with genes that were differentially expressed in nurses and foragers [49]. Similarly, we found a significant degree of overlap (hypergeometric test, P<0.05) between DEGs associated with BP exposure in our study and BP-related DEGs identified in [37] after 15 days of continuous exposure. Thus, long-term changes in gene expression associated with impacts of BP exposure on the transition from nursing to foraging tasks overlap significantly with short-term changes in brain expression patterns associated with the stimulation of foraging behavior by BP. This ultimately suggests that behavioral plasticity to utilize common suites of genes at vastly different time scales seems.

**Table 5:** Overlaps between pheromone-related genes and those of Alaux et al.

|  | Genes represented in both | BP genes | Alaux et al., BP after 5 days | Alaux et al., BP after 15 days | Overlap BP5 | Overlap BP15 |
| --- | --- | --- | --- | --- | --- | --- |
| BP vs Control | 6039 | 49 | 104 | 85 | 1 | 2* |
| *significantly greater overlap of genes than expected by chance; P<0.05; hypergeometric test |  |  |  |  |  |  |

**Table 6:** Overlaps between foraging-related genes and Whitfield et al.

|  | Genes represented in both | Foraging-related | Whitfield et al | Overlapping genes |
| --- | --- | --- | --- | --- |
| BP vs Control | 6039 | 264 | 839 | 48* |
| *significantly greater overlap of genes than expected by chance; P<0.05; hypergeometric test |  |  |  |  |

### Weighted gene co-expression network analysis

We used WGCNA to construct networks of genes based solely on the similarity of their expression patterns to organize co-expressed genes into groups, called modules. These modules were constructed independently of trait information and were then correlated to traits using a generalized linear model. Specifically, we looked at relationships between each module and three traits of interest: pollen vs. nectar foraging, BP vs. control, and EBO vs. control. In this way, the WGCNA identified 16 modules that were significantly correlated to forager type, exposure to BP, exposure to EBO, or a combination thereof (GLM, P<0.05; Fig. 4). Fourteen modules were significantly correlated with only one trait. Module 10 was the only module that was associated with all traits, while Module light cyan was associated with forager type and EBO exposure, but not BP exposure. For each module, the most highly connected gene in the network was identified (Table 7), providing a list of candidate genes. The top five most connected genes for each module can be found in the Supplementary Materials.

**Table 7:**
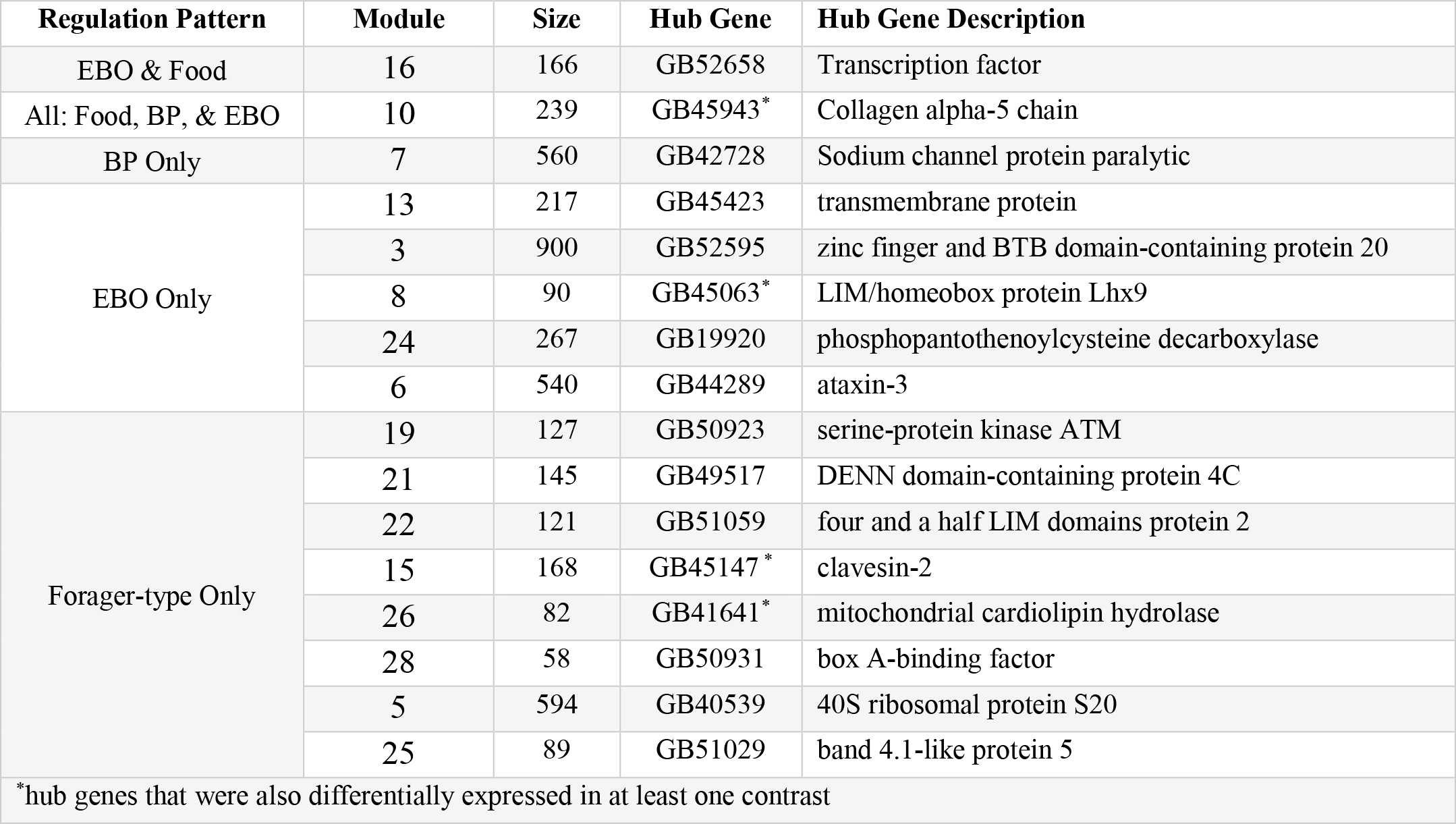
WGCNA Module Hub genes

**Figure 4.**
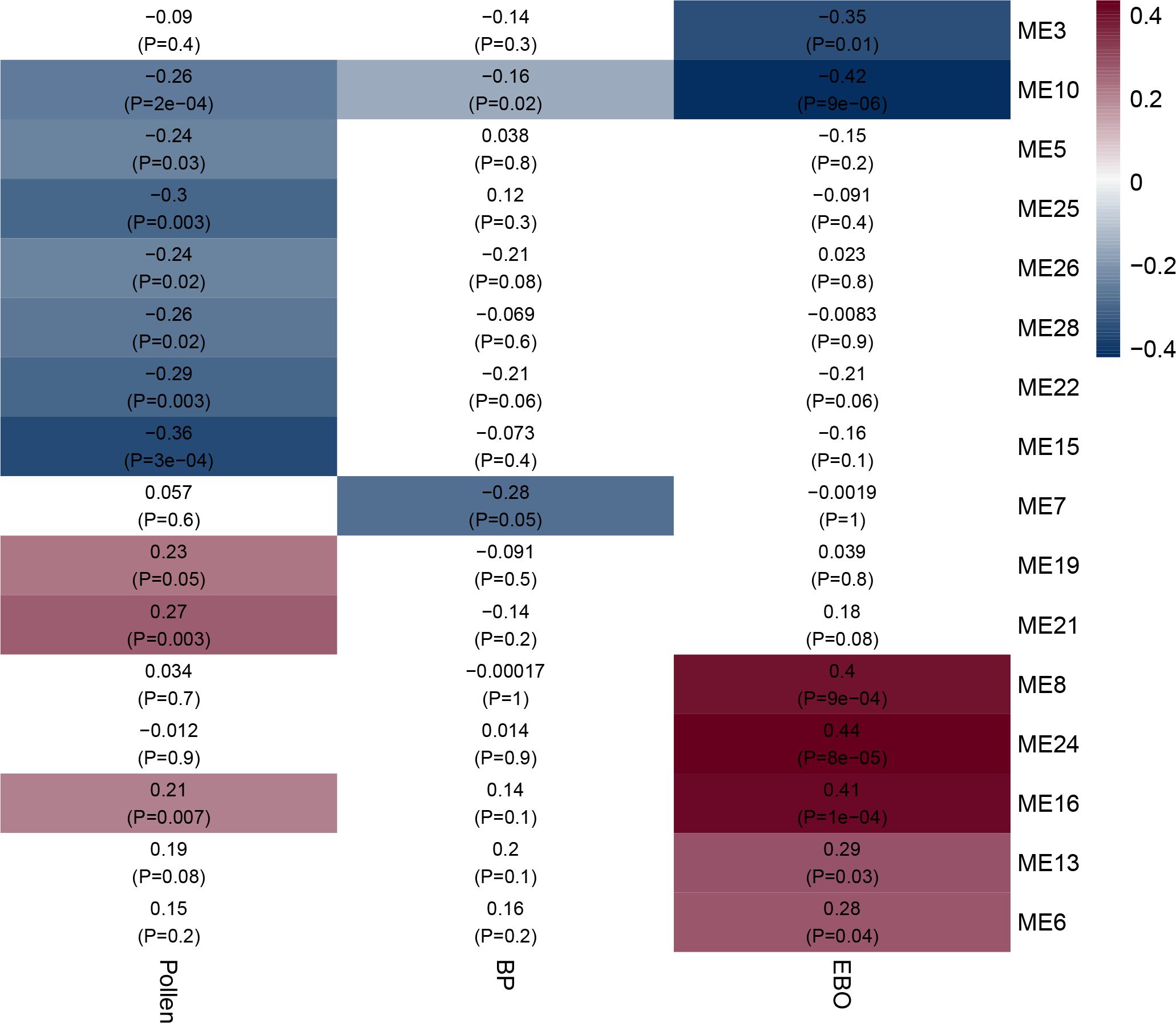
Weighted gene co-expression network analysis. Rows represent gene modules. Columns represent sample traits. Each cell contains two values: a correlation coefficient between the module and sample trait and the associated p-value in parentheses. Significant correlations are colorized according correlation coefficient, varying from high values in yellow to low values in purple.

To better understand the functions of the gene modules identified in this analysis, we performed KEGG (Kyoto Encyclopedia of Genes and Genomes) pathway analysis on three modules (Table 8). Module 10 was chosen based on its significant correlation with food and both brood pheromones, and modules 3 and 7 were selected based on their strong correlations with BP and EBO, respectively. Module 10 was enriched for KEGG pathways related to metabolic pathways, carbon metabolism, fatty acid metabolism, and peroxisomes (Wilcoxon, P<0.05). Module 7 was significantly enriched for glycerophospholipid metabolism, neuroactive ligand-receptor interaction, and hippo signaling pathway (Wilcoxon, P<0.05). Module 3 was enriched for metabolic pathways like FoxO and AGE-RAGE signaling pathways, development pathways like wnt signaling, and immune pathways like Toll and lmd signaling pathways (Wilcoxon, P<0.05).

**Table 8:** KEGG analysis of selected WGCNA modules

| Module | Trait association | Significantly enriched KEGG pathways (P<0.05) | Significantly enriched GO categories (EASE <0.05) |
| --- | --- | --- | --- |
| 10 | BP, EBO, Forager-type | Metabolic pathways<br>Carbon metabolism<br>Fatty acid metabolism<br>Peroxisome | Integral components of membrane,<br>Fatty acid biosynthetic process |
| 7 | BP alone | Glycerophospholipid metabolism<br>Neuroactive ligand-receptor interaction<br>Hippo signaling pathway | Ion channel |
| 3 | EBO alone | Pentose and glucuronate interconversions<br>Metabolic pathways<br>FOXO signaling pathway<br>Neuroactive ligand-receptor interaction<br>Lysosome<br>Wnt signaling pathway<br>Dorso-ventral axis formation<br>Notch signaling pathway<br>Toll and Imd signaling pathway<br>AGE-RAGE signaling pathway in diabetic complications | Integral components of membrane |

## Discussion

In the present study, we investigated the genes and transcriptional pathways underlying rapid behavioral responses to pheromone signals in honey bee foragers specializing on pollen or nectar foraging. We hypothesized that two larval pheromones, brood pheromone (BP) and E-beta-ocimene (EBO), would regulate a common set of foraging genes in the brain, and that they would affect gene expression more strongly in pollen foragers than nectar foragers. We found that nectar and pollen foragers have distinguishable gene expression profiles, and that both larval pheromones do indeed regulate a shared set of genes and transcriptional pathways; however, these transcriptional pathways more strongly affect nectar foragers than pollen foragers. Moreover, comparisons with previous studies suggest that similar genes regulate the ontogeny of foraging behavior and foraging task specialization, and a common set of genes mediates both short- and long-term responses to BP. Thus, our study identified mechanistic links between larval pheromone communication and foraging specialization and suggests that common transcriptional pathways may regulate behavior across time scales.

Pheromone signals can cause various changes in the receiver’s behavior depending on the behavioral and physiological state of the individual, sometimes with consequences for survival and reproduction [50, 51]. Honey bee larval pheromones are particularly interesting because they trigger similar behavioral changes on two timescales: long-term behavioral transition from nursing to foraging and short-term upregulation of pollen foraging behavior. Previous studies have examined the genetic and behavioral differences associated with preference for nectar versus pollen foraging [28, 52–55]. Furthermore, larval pheromones elicit specific response in pollen foragers. For example, exposure to brood pheromone (BP) increased colony-level pollen foraging 2.5 fold [44], as well as the ratio of pollen to non-pollen foragers [45], and the individual effort of pollen foragers [45]. However, previous to this study, there were no documented impacts of exposure to brood pheromones on nectar foraging.

The present study demonstrates for the first time that there are also transcriptional differences between nectar and pollen foragers in the mushroom bodies of honey bees. Several quantitative trait loci have been identified which underlie colony-level variation in the propensity to collect pollen versus nectar, and these loci are associated with variation in concentration of nectar collected and the amount of pollen and nectar brought back to the hive [54, 56]. In our study, foraging specialization on nectar versus pollen foraging was associated with substantial differences in gene expression profiles (with almost 400 DEGs; Table 1), and with variation among nectar and pollen foragers, which accounted for 46% percent of the overall variation in DEGs (Fig. 3). To elucidate transcriptional pathways that respond to larval pheromones, we utilized weighted gene correlation network analysis (WGNCA) to provide a more detailed view of molecular processes associated with traits of interest [57, 58]. WGNCA identified 16 genetic modules that were significantly correlated with foraging or pheromone exposure (Fig. 4), most of which were associated with foraging specialization (Fig. 4).

Short exposure to both BP and EBO significantly altered gene expression profiles in the brains of foragers, and both pheromones regulated overlapping sets of genes. Exposure to EBO was associated with 169 DEGs, which was nearly three times greater than the number of DEGs regulated by BP (Table 1). Yet, even in this limited gene set, there was a statistically significant overlap in the DEGs regulated by BP and EBO (Table 2), and the overlapping genes were enriched for fatty acid metabolism. Hierarchical clustering and principal component analyses confirmed that pheromone exposure had strong and consistent effects on gene expression profiles. Furthermore, WGCNA revealed that module 10, representing 239 genes with correlated expression patterns, was significantly downregulated in samples exposed to either pheromone. Together, these results suggest that BP and EBO regulate overlapping genetic modules and pathways that are enriched for energy metabolism. Decreasing whole-brain energy metabolism, including that of fatty-acids, is associated with long-term behavioral transition from in-hive tasks to foraging tasks, suggesting that larval pheromones regulate foraging behavior by specifically activating pathways involved the natural ontogeny for foraging behavior [59].

Our data supported our hypothesis that exposure to larval pheromones alters expression of foraging related genes, but contrary to our predictions, the pheromones had more pronounced effects on gene expression in nectar foragers than pollen foragers. There was a common set of DEGs that were associated with both pheromone treatment and foraging specialization (Table 3), which was driven primarily by DEGs in nectar foragers but not pollen foragers (Table 4). Hierarchical clustering analysis show that, for the most part, samples were clustered into pollen and nectar foraging “branches,” with the intriguing exception that nectar foragers exposed to EBO had expression profiles that were more like those of pollen foragers (Table 4). Similarly, principal components analysis show that nectar foragers exposed to EBO cluster more closely with pollen foragers than other nectar foragers (Fig 3). The gene network analysis revealed that two modules were associated with both pheromone treatment and foraging, one of which was enriched for membrane components and energy metabolism (Table 8). These results suggest that one mechanism by which larval pheromones modulate colony-level pollen foraging behavior is by downregulating metabolic pathways in the brains of nectar foragers, which is consistent with the role that energy metabolism plays in the ontogeny of foraging behavior [59]. Pankiw et al. [44] found that short exposure to BP increased pollen foraging, but did not observe task-switching of nectar foragers to pollen foraging, which the authors found puzzling. Our results indicate that one explanation may be that even after short exposures to larval pheromones, nectar foragers may be primed to switch to pollen foraging even before they actually make the behavioral transition, which may be a way to buffer against ephemeral swings in the nutritional demands of developing larvae.

Changes in the same behavior at different time scales, such as ontogeny of pollen foraging and pheromonal upregulation of pollen foraging, may utilize similar molecular mechanisms. We reached this intriguing conclusion after comparing our results to those of two landmark honey bee transcriptome studies [37, 49]. Whitfield et al. [49] compared nurses and foragers, controlling for age, and found over 1,000 DEGs. Alaux et al. [37] was the first to study the effects of brood pheromone on gene expression, and found more than 200 DEGs between age-matched bees that were exposed to BP continuously for multiple days (i.e., 5 or 15 days) and those that were not exposed. To test the degree of overlap between our results and those from previous studies, we compared 1) the number of DEG between nectar and pollen foragers in our study with those identified by Whitfield et al. [62], and 2) the DEGs between pheromone treatments in our study with those identified by Alaux et al. [37]. We found significant overlaps between DEGs identified in the present study and those of Whitfield et al. (P<0.001) and of Alaux et al. (P<0.001). The significant overlap between our study and the two microarray studies, which validate the expression patterns related to foraging specialization and brood pheromone exposure, suggests that foraging-related gene expression shows a degree of consistenty across time scales (see [58]), and supports the idea that pheromones regulate the transcriptional pathways underlying foraging specialization.

The results of our study lay the groundwork for several intriguing lines of inquiry for future studies. First, exposing foragers to a short pulse of BP, which stimulates immediate foraging, regulates a similar set of genes as bees as exposing nurses to BP for 5 days, which modulates the transition from in-hive tasks to foraging. BP potentially regulates foraging behavior, at least in part, by priming their to receptivity to foraging-related or social stimuli, even before nurse bees have made the physiological transition to foraging tasks. This could conceivably involve genes implicated in both foraging and division of labor (e.g., *Malvolio*, a manganese transporter) [60], or neurochemical regulatory pathways involving octopomine, which has shown to modulate responsiveness to both foraging-stimuli and to BP [61, 62]. Furthermore, our results suggest the hypothesis that social pheromones upregulate pollen foraging by decreasing the expression of nectar foraging genes in the brain, and this would also be productive line of inquiry for future studies. Lastly, our data suggest that rapid changes in brain gene expression in nectar foragers may happen prior to task-switching to buffer against ephemeral environmental conditions. Short-term exposure to larval pheromones may “prime” nectar foragers to switch preferences to pollen foraging, and this switch could occur under conditions of prolonged exposure to brood pheromone. Thus, our study provides a framework for hypothesis-driven experiments examining the impacts of pheromone exposure on task specialization and division of labor.

DEGs and WGCNA modules related to both pheromone treatment and foraging specialization were enriched for several metabolic pathways, including fatty-acid metabolism, which suggests that metabolic processes and lipid signaling in integration centers of the honey bee brain may play a role in behavioral plasticity. The transition from nursing to foraging involves large-scale changes in metabolic pathways, including reductions in lipid stores and changes in insulin signaling [63]. These physiological changes during the transition from in-hive tasks to foraging are associated with changes in energy metabolism (including insulin signaling), gustatory response, and foraging preferences for nectar vs pollen [64, 65]. Therefore, the prominence of energy metabolism, lipid signaling pathways, and related metabolic pathways in our brain transcriptome data supports the idea that these pathways in the brain play a role in insect behavior [66, 67]. Other studies have demonstrated the importance of brain metabolic processes on influencing individual variation in behavior, particularly aggression [67–69]. The enrichment of metabolic pathways in DEGs and the prominence of FOXO signaling pathway in our gene co-expression networks further supports the role of insulin signaling pathways in mediating neuronal function and behavior in insects [66, 67]. For example, an insulin binding protein, *Queen brain-selective protein-1 (Qbp-1)*, was differentially expressed in response to pheromone treatment and is related to FOXO signaling. Module 3 was enriched for FOXO signaling and significantly correlated with EBO treatment, so its hub genes may serve as useful candidate genes for subsequent studies investigating the impact of insulin signaling on pheromone communication and foraging.

## Conclusions

The neural circuits and molecular pathways underlying behavioral plasticity and task specialization are complex, and our study demonstrates that foraging behavior is regulated in part by common suites of genes across time scales, from long-term behavioral plasticity (nurse to forager) to individual variation in task specialization (pollen vs nectar). Our study further confirms that pheromone communication has a profound effect on gene expression within hours of exposure, and more importantly, that social signals (i.e. pheromones) may invoke foraging-related transcriptional pathways to upregulate pollen foraging at both long and short-time time scales. Moreover, there is clear interaction between individual variation in task specialization and responses to social signals (i.e. pheromones), and these social signals seem to be manipulating brain energy metabolism to elicit foraging behavior. Because the mechanisms underlying foraging behavior are deeply conserved in animals, a detailed mechanistic understanding of the foraging in honey bees may provide insights into mechanisms involved in division of labor in insect societies, foraging preference in animals, and complex behavioral phenotypes in general.

## Methods

### Animals and Experimental Design

We created single-cohort colonies (using same-aged workers) from a common source colony with a naturally mated queen to avoid differences in behavior and gene expression due to variation in age and genetic background; thus, all bees used in the study were half-sisters. Because workers performing any given task in a natural colony can vary widely in age, we constructed single cohort colonies using workers that emerged as adults within a 48-hr period, minimizing differences in age and experience among individuals. After one week, some of the bees in single-cohort colonies transition quickly to foraging (Robinson et al. 1989).

Three such colonies—each provided with an identical starting population (1,000 bees), honey and pollen resources—were placed in a large outdoor enclosure (approximately 20’ x 50’ft) at the Texas A & M University Riverside Campus. Each colony was provided with two frames: one frame laden with pollen and honey stores, and one empty frame. In addition to frames full of honey and pollen inside the colony, feeders full of 50% (w/v) sucrose solution and fresh pollen (collected from pollen traps on unrelated honey bee colonies) were placed in front of each colony daily, and bees that foraged on each resource were given different color dots of enamel paint (Testors, Rockford, IL) on the thorax based on their preference for pollen or nectar feeders. Only bees with multiple paint marks of a single color were used in this study because multiple same-color marks demonstrated consistent preference for pollen or nectar, respectively. Each colony was also provided a strip of queen pheromone (PseudoQueen, Contech Industries, Victoria, BC, Canada) to prevent colonies from developing a “queenless” physiological state and to control for the variation in queen quality that would inherently occur when using live queens. Colonies did not receive any frames containing brood. Although broodless colonies are not the default state of a colony, it is nevertheless a natural occurrence when queens die for any number of reasons [52]. The absence of brood controlled for the natural variation in brood pheromones that may have occurred with the presence of real brood, and minimized the amount of beekeeping interference necessary to maintain identical colony conditions.

After two weeks, colonies were exposed to field-relevant dosages (5,000 larval equivalents) of EBO, BP, or a paraffin oil control. We used a BP blend characteristic of older larvae that has been shown to strongly upregulate pollen foraging, as done previously by Ma et al. [45]. Hive entrances were blocked during the one-hour period during which the pheromone treatment was applied, and any foragers outside the colony during that time were removed from the experiment when they landed on the blocked entrance. When the entrances were opened, forager bees (previously marked as nectar or pollen foragers, as described above) were collected as they landed on a pollen or nectar feeder, but before they initiated feeding. Six pollen foragers and six nectar foragers were collected from each colony and placed immediately in dry ice for later brain dissection. Subsequently, we pooled pairs of bees to generate the RNA samples. Thus, in total, there were three pollen forager and three nectar forager samples for each of the three colonies representing the control, BP and EBO treatments (Fig. 1; total number of samples = 18). Sampled individuals were stored at −80°C until they were dissected.

### Brain Dissection

Insect mushroom bodies are considered important integration centers of the brain because of their role in multimodal information processing and their association with learning and memory [70–72]. These factors make mushroom bodies an ideal candidate brain region to investigate temporal dynamics of communication and behavior. Therefore, mushroom bodies of the brain were dissected from sampled individuals by placing them on dry ice to prevent thawing and degradation of transcripts, as in [37, 40]. However, in our study, the brains were not freeze-dried to facilitate dissection of the mushroom bodies. For each sample, RNA from two brains were extracted using the RNeasy Mini Kit (Qiagen, Valencia, CA) following the manufacturer’s protocol, and pooled RNA quantity and quality were assayed using a Qubit Fluorometer (Invitrogen, Carlsbad, CA). cDNA library preparation and sequencing were performed by the Genomic Sequencing and Analysis Facility at the University of Texas at Austin using an Illumina HiSeq 4000 (Illumina, San Diego, CA) sequencer. All 18 samples were barcoded and split across four lanes to control for sequencing bias. A total of 18 RNA-seq single-end 50bp libraries were generated, with three libraries for each treatment group from each colony (Fig 1).

### Data Analysis

Reads were trimmed using Trimmomatic [73] to remove adapter sequences, low-quality reads, and short reads (<36 bp). Kallisto software was used to build a transcriptome index based on a recently updated honey bee reference genome annotation (Amel_HAv3.1), and subsequently to quantify the abundance of transcripts represented in each sample [74]. The R package tximport was then used to import transcript abundances and generate a gene-level count matrix that was scaled to both library size and transcript length [75]. The transcript-gene correspondence was derived from the genome annotation using the R package rtracklayer [76]. The R package DESeq2 [77] was used to collapse technical replicates (i.e. identical samples across multiple sequencing lanes) and perform differential gene expression analysis with pheromone treatment, forager type, and the interaction of pheromone treatment and forager type as fixed effects in a generalized linear model. Only genes with an abundance of at least one transcript per million in all samples were used in the analysis. Genes whose expression differed between groups were considered differentially expressed when they had a false discovery rate (FDR) of <0.05. Gene ontology analysis was performed using the Database for Annotation, Visualization and Integrated Discovery (DAVID v6.8) to better understand the biological relevance of differentially expressed genes (DEGs) [78].

The expression patterns of DEGs were further analyzed by performing unsupervised hierarchical clustering (Fig. 2; Supplementary Fig 1) and PCA (Fig. 3) on gene expression data normalized through variance stabilizing transformation. Hierarchical clustering was performed in R and visualized using the pheatmaps package [79]. Genes were clustered using the Ward method and samples were clustered based on manhattan distance. Principal component analysis (PCA) was performed to find the linear combinations of genes that explained the maximum amount of variation in the data, producing a series of orthogonal factors (i.e. principal components). PCA results were visualized in ggplot2 [80].

### Gene co-expression network analysis

To generate a global unsupervised overview of the gene expression data, we utilized weighted gene correlation network analysis [57] to identify groups of co-expressed genes and assess the relationship between these groups and experimental treatments. WGCNA constructs networks of genes based solely on the similarity of their expression patterns and organized them into groups of co-expressed genes, called modules. Module assignment is unsupervised and independent of sample trait information (e.g., pheromone treatment, forager-type), and subsequently, these gene modules can be correlated with traits of interests. In this way, WGCNA can supplement other genomic and bioinformatic methods to provide a more detailed view of molecular processes associated with traits of interest.

Variance stabilized gene expression data were grouped into modules based on similarity of expression patterns. Because genes within each module showed very highly correlated patterns, the first principal component of the genes within a module was used to represent the entire module (module ‘eigengene’). Then, these module representatives were correlated with sample traits using a generalized linear model, with forager-type and pheromone as fixed effects (Fig. 4). Minimum module size was set to 30, and deep split was set to 2. Modules were built with a standardized connectivity score of −2.5, and module definition was based on “hybrid” branch cutting. A signed gene co-expression network was constructed with a soft threshold of 10. Modules were merged based on a cut height of 0.1. Module eigengenes were correlated with sample traits using a generalized linear model with forager-type, pheromone exposure treatment, and their interaction as fixed effects.

### Overlap of differentially expressed genes with previous studies

Hypergeometric tests were used to assess whether there was a significant overlap of differentially expressed genes when compared to other studies. Specifically, we tested overlap with genes regulated by long-term exposure to brood pheromone [37] and genes that varied between nurses and foragers [49]. These two studies utilized microarrays containing approximately 5,500 genes identified in an earlier genome assembly version. For consistency, microarray probes were mapped to current official honey bee gene set, as done in Khamis et al. [81]. The degree of overlap between our data and data from these two studies were assessed using hypergeometric tests in the base stats package of R.

## LIST OF ABBREVIATIONS

BP: Brood Pheromone
DEG: Differentially expressed gene
EBO: E-beta-ocimene
FDR: False discovery rate
GO: Gene ontology
KEGG: Kyoto Encyclopedia of Genes and Genomes
PC: Principal component
PCA: Principal components analysis
RNA: Ribonucleic acid
RNA-Seq: RNA sequencing
WGCNA: Weighted gene co-expression network analysis

## DECLARATIONS

### Ethics approval and consent to participate

Not applicable

### Consent for publication

Not applicable

### Availability of data and material

The raw RNA-Seq data were deposited in the NCBI Sequence Read Archive under submission number SUB5286697 and BioProject number PRJNA528102: http://www.ncbi.nlm.nih.gov/bioproject/528102.

### Author’s contributions

RM conceived and performed the experiment. RM, JR, and CMG designed the experiment. RM and CMG analyzed the data. All authors participated in writing the manuscript.

## Acknowledgements

We would like to thank ET Ash and LA Ward for beekeeping assistance; C Stengl for generous support of the graduate program at UT Austin; and LE Gilbert, HA Hofmann, S Jha, UG Mueller, and members of the Grozinger lab for comments on the manuscript.

## Supplementary Materials

Table and Figure Legends

Supplementary Table 1. Transcriptome assembly quality metrics averaged across four technical replicates per sample, given in numbers of sequences per sample and as percentages of original sequencing reads per sample.

Supplementary Table 2. Entrez gene IDs of differentially up-regulated and down regulated genes in the context of pheromone exposure, foraging task-specialization, and their interaction.

Supplementary Table 3. Dictionary of transcripts, Entrez Gene ID, BeeBase ID, and Accession numbers for all transcripts in the study.

Supplementary Figure 1. Hierarchical clustering with multiscale bootstrap resampling confirms that bees exposed to identical pheromone exposure and forager-type produced distinctive transcriptional profiles in honey bee brains. For each cluster, two p-values are displayed on edges, expressed as percentages. The red number on the left represents the Approximately Unbiased (AU) method, and the green number on the right represents bootstrap probability (BP). Red rectangles indicate significant clusters with AU values greater than 95, indicating strongly supported clusters. Samples names denote pheromone exposure (i.e. Control (X), brood pheromone (BP), and E-beta-ocimene (EBO), forager type (Pollen (pol) vs nectar (N), or and sample number (1-3). This analysis used all 533 DEGs identified in this study.

Supplementary Figure 2. Clustering of variance stabilized gene expression data during co-expression network analysis. Modules were formed independently of sample information, and the colors under the cluster dendrogram indicate the assignment of co-expressed genes to modules. “Dynamic tree cut” colors indicate original module assignments before merging similar modules (cut height 0.1), while “Merged dynamic” colors represent final module assignments after merging similar modules.

## References

1. Seeley T. The wisdom of the hive: the social physiology of honey bee colonies. 2009.

2. Michener CD. Comparative social behavior of bees. Annu Rev Entomol. 1969;14:299–342.

3. Simpson C. The evolutionary history of division of labour. Proc R Soc B Biol Sci. 2012;279:116–21.

4. Robinson GE, Fernald RD, Clayton DF. Genes and social behavior. Science. 2008;322:896–900.

5. Amador-Vargas S, Gronenberg W, Wcislo WT, Mueller U. Specialization and group size: brain and behavioural correlates of colony size in ants lacking morphological castes. Proceedings Biol Sci. 2015;282:20142502.

6. Rueffler C, Hermisson J, Wagner GP. Evolution of functional specialization and division of labor. Proc Natl Acad Sci. 2011.

7. Robinson GE, Barron AB. Epigenetics and the evolution of instincts. Science. 2017;356:26–7.

8. Barta Z. Individual variation behind the evolution of cooperation. Philos Trans R Soc Lond B Biol Sci. 2016;371:20150087.

9. NESCent Working Group on Integrative Models of Vertebrate Sociality: Evolution, Mechanisms and EP, Hofmann HA, Beery AK, Blumstein DT, Couzin ID, Earley RL, et al. An evolutionary framework for studying mechanisms of social behavior. Trends Ecol Evol. 2014;29:581–9.

10. Ben-Shahar Y, Robichon A, Sokolowski MB, Robinson GE. Influence of Gene Action Across Different Time Scales on Behavior. Science (80-). 2002;296:741–4.

11. Ortíz-Barrientos D, Noor M a F. Evidence for a one-allele assortative mating locus. Science (80-). 2005;310:1467.

12. Elsik CG, Worley KC, Bennett AK, Beye M, Camara F, Childers CP, et al. Finding the missing honey bee genes: lessons learned from a genome upgrade. BMC Genomics. 2014;15:86.

13. Aubin-Horth N, Renn SCP. Genomic reaction norms: Using integrative biology to understand molecular mechanisms of phenotypic plasticity. Mol Ecol. 2009;18:3763–80.

14. Insel TR. The Challenge of Translation in Social Neuroscience: A Review of Oxytocin, Vasopressin, and Affiliative Behavior. Neuron. 2010;65:768–79.

15. Barron AB, Maleszka J, Vander Meer RK, Robinson GE, Maleszka R. Comparing injection, feeding and topical application methods for treatment of honeybees with octopamine. J Insect Physiol. 2007;53:187–94.

16. Goodson JL. The vertebrate social behavior network: Evolutionary themes and variations. Horm Behav. 2005;48:11–22.

17. Insel TR, Young LJ. Neuropeptides and the evolution of social behavior. Curr Opin Neurobiol. 2000;10:784–9.

18. Donaldson ZR, Young LJ. Oxytocin, vasopressin, and the neurogenetics of sociality. Science. 2008;322:900–4.

19. Young RL, Ferkin MH, Ockendon-Powell NF, Orr VN, Phelps SM, Pogány Á, et al. Conserved transcriptomic profiles underpin monogamy across vertebrates. Proc Natl Acad Sci. 2019;:201813775.

20. Toth AL, Varala K, Newman TC, Miguez FE, Hutchison SK, Willoughby DA, et al. Wasp gene expression supports an evolutionary link between maternal behavior and eusociality. Science. 2007;318:441–4.

21. Hau M, Goymann W. Endocrine mechanisms, behavioral phenotypes and plasticity: known relationships and open questions. Front Zool. 2015;12 Suppl 1:S7.

22. Schaefer N, Rotermund C, Blumrich E-M, Lourenco M V., Joshi P, Hegemann RU, et al. The malleable brain: plasticity of neural circuits and behavior - a review from students to students. J Neurochem. 2017;142:790–811.

23. Rittschof CC, Hughes KA. Advancing behavioural genomics by considering timescale. Nat Commun. 2018;9:489.

24. Weir J. Polyethism in workers of the antMyrmica. Insectes Soc. 1958;5:97–128.

25. Robinson GE. Regulation of honey bee age polyethism by juvenile hormone. Behav Ecol Sociobiol. 1987;20:329–38.

26. Calderone NW, Page RE. Genotypic variability in age polyethism and task specialization in the honey bee, Apis mellifera (Hymenoptera: Apidae). Behav Ecol Sociobiol. 1988;22:17–25.

27. Seeley TD. Honeybee ecology: a study of adaptation in social life. Princeton, New Jersey: Princeton University Press; 2014.

28. Fewell JH, Page Jr. RE. Genotypic Variation in Foraging Responses to Environemental Stimuli by Honey Bees, Apis mellifera. Experientia. 1993;49:1106–12.

29. Haydak MH. Honey Bee Nutrition. Annu Rev Entomol. 1970;15:143–56.

30. Camazine S. The regulation of pollen foraging by honey bees: how foragers assess the colony’s need for pollen. Behav Ecol Sociobiol. 1993;32:265–72.

31. Pankiw T, Page Jr RE. Response thresholds to sucrose predict foraging division of labor in honeybees. Behav Ecol Sociobiol. 2000;47:265–7.

32. Kocher SD, Ayroles JF, Stone EA, Grozinger CM. Individual variation in pheromone response correlates with reproductive traits and brain gene expression in worker honey bees. PLoS One. 2010;5.

33. Metz BN, Pankiw T, Tichy SE, Aronstein KA, Crewe RM. Variation in and responses to brood pheromone of the honey bee (*Apis mellifera L*.). J Chem Ecol. 2010;36:432–40.

34. Grozinger CM, Sharabash NM, Whitfield CW, Robinson GE. Pheromone-mediated gene expression in the honey bee brain. Proc Natl Acad Sci U S A. 2003;100 suppl. 2:14519–25.

35. Alaux C, Robinson GE. Alarm pheromone induces immediate-early gene expression and slow behavioral response in honey bees. J Chem Ecol. 2007;33:1346–50.

36. Alaux C, Sinha S, Hasadsri L, Hunt GJ, Guzmán-Novoa E, DeGrandi-Hoffman G, et al. Honey bee aggression supports a link between gene regulation and behavioral evolution. Proc Natl Acad Sci U S A. 2009;106:15400–5.

37. Alaux C, Le Conte Y, Adams H, Rodriguez-Zas S, Grozinger C, Sinha S, et al. Regulation of brain gene expression in honey bees by brood pheromone. Genes Brain Behav. 2009;8:309–19.

38. Slessor KN, Winston ML, Le Conte Y. Pheromone communication in the honeybee (*Apis mellifera L*.). J Chem Ecol. 2005;31:2731–45.

39. Whitfield CW, Cziko A-M, Robinson GE. Gene expression profiles in the brain predict behavior in individual honey bees. Science. 2003;302:296–9.

40. Grozinger CM, Sharabash NM, Whitfield CW, Robinson GE. Pheromone-mediated gene expression in the honey bee brain. Proc Natl Acad Sci U S A. 2003;100 Suppl 2 suppl 2:14519–25.

41. Lutz CC, Robinson GE. Activity-dependent gene expression in honey bee mushroom bodies in response to orientation flight. J Exp Biol. 2013;216 Pt 11:2031–8.

42. Wanner KW, Nichols AS, Walden KKO, Brockmann A, Luetje CW, Robertson HM. A honey bee odorant receptor for the queen substance 9-oxo-2-decenoic acid. Proc Natl Acad Sci U S A. 2007;104:14383–8.

43. Beggs KT, Glendining KA, Marechal NM, Vergoz V, Nakamura I, Slessor KN, et al. Queen pheromone modulates brain dopamine function in worker honey bees.

44. Pankiw T, Page Jr RE, Kim Fondrk M. Brood pheromone stimulates pollen foraging in honey bees (*Apis mellifera*). Behav Ecol Sociobiol. 1998;44:193–8.

45. Pankiw T. Brood Pheromone Regulates Foraging Activity of Honey Bees (Hymenoptera: Apidae). J Econ Entomol. 2004;97:748–51.

46. Traynor KS, Le Conte Y, Page REJ. Age matters: Pheromone profiles of larvae differentially influence foraging behaviour in the honeybee, *Apis mellifera*. Anim Behav. 2015;99:1–8.

47. Ma R, Mueller UG, Rangel J. Assessing the role of β-ocimene in regulating foraging behavior of the honey bee, *Apis mellifera*. Apidologie. 2016;47:135–44.

48. Ma R, Villar G, Grozinger CM, Rangel J. Larval pheromones act as colony-wide regulators of collective foraging behavior in honeybees. Behav Ecol. 2018;29:1132–41.

49. Whitfield CW. Gene Expression Profiles in the Brain Predict Behavior in Individual Honey Bees. Science (80-). 2003;302:296–9.

50. Le Conte Y, Hefetz A. Primer pheromones in social hymenoptera. Annu Rev Entomol. 2008;53:523–42.

51. Wyatt T. Pheromones and Animal Behaviour: communication by smell and taste. 1st Editio. Cambridge: Cambridge University Press; 2003.

52. Winston ML. The Biology of the Honey Bee. Cambridge: Harvard University Press; 1991.

53. Fewell J, Winston M. Colony state and regulation of pollen foraging in the honey bee, Apis mellifera L. Behav Ecol Sociobiol. 1992;30:387–93.

54. Robinson GE, Page RE. Genetic determination of nectar foraging, pollen foraging, and nest-site scouting in honey bee colonies. Behav Ecol Sociobiol. 1989;24:317–23.

55. Sagili RR, Pankiw T, Metz BN. Division of labor associated with brood rearing in the honey bee: how does it translate to colony fitness? PLoS One. 2011;6:e16785.

56. Ruppell O, Pankiw T, Page RE. Pleiotropy, epistasis and new QTL: The genetic architecture of honey bee foraging behavior. J Hered. 2004;95:481–91.

57. Langfelder P, Horvath S. WGCNA: an R package for weighted correlation network analysis. BMC Bioinformatics. 2008;9:559.

58. Lutz CC, Rodriguez-Zas SL, Fahrbach SESE, Robinson GE. Transcriptional response to foraging experience in the honey bee mushroom bodies. Dev Neurobiol. 2012;72:153–66.

59. Ament SA, Corona M, Pollock HS, Robinson GE. Insulin signaling is involved in the regulation of worker division of labor in honey bee colonies. 2008.

60. Greenberg JK, Xia J, Zhou X, Thatcher SR, Gu X, Ament S a, et al. Behavioral plasticity in honey bees is associated with differences in brain microRNA transcriptome. Genes Brain Behav. 2012;11:660–70.

61. A. B, D. S, G. R. Octopamine modulates responsiveness to foraging-related stimuli in honey bees (Apis mellifera). J Comp Physiol A Sensory, Neural, Behav Physiol. 2002;188:603–10.

62. Schulz DJ, Barron AB, Robinson GE. A Role for Octopamine in Honey Bee Division of Labor. Brain Behav Evol. 2002;60:350–9.

63. Ament SA, Chan QW, Wheeler MM, Nixon SE, Johnson SP, Rodriguez-Zas SL, et al. Mechanisms of stable lipid loss in a social insect. J Exp Biol. 2011;214 Pt 22:3808–21.

64. Wang Y, Mutti NS, Ihle KE, Siegel A, Dolezal AG, Kaftanoglu O, et al. Down-Regulation of Honey Bee IRS Gene Biases Behavior toward Food Rich in Protein. PLoS Genet. 2010;6:e1000896.

65. Wang Y, Brent CS, Fennern E, Amdam G V. Gustatory Perception and Fat Body Energy Metabolism Are Jointly Affected by Vitellogenin and Juvenile Hormone in Honey Bees. PLoS Genet. 2012;8:e1002779.

66. Wu Q, Brown MR. Signaling and Function of Insulin-Like Peptides in Insects. Annu Rev Entomol. 2005;51:1–24.

67. Rittschof CC, Grozinger CM, Robinson GE. The energetic basis of behavior: Bridging behavioral ecology and neuroscience. Curr Opin Behav Sci. 2015;6:19–27.

68. Chandrasekaran S, Rittschof CC, Djukovic D, Gu H, Raftery D, Price ND, et al. Aggression is associated with aerobic glycolysis in the honey bee brain ^1^. Genes, Brain Behav. 2015;14:158–66.

69. Li-Byarlay H, Rittschof CC, Massey JH, Pittendrigh BR, Robinson GE. Socially responsive effects of brain oxidative metabolism on aggression. PNAS. 2014;111:12533–7.

70. Menzel R. The insect mushroom body, an experience-dependent recoding device. J Physiol. 2014;108:84–95.

71. Farris SM. Are mushroom bodies cerebellum-like structures? Arthropod Struct Dev. 2011;40:368–79.

72. Lihoreau M, Latty T, Chittka L. An exploration of the social brain hypothesis in insects. 2012;3 November:1–7.

73. Bolger A, Lohse M, Usadel B. Trimmomatic: a flexible trimmer for Illumina sequence data. Bioinformatics. 2014.

74. Bray NL, Pimentel H, Melsted P, Pachter L. Near-optimal probabilistic RNA-seq quantification. Nat Biotechnol. 2016;34:525–7.

75. Soneson C, Love MI, Robinson MD. Differential analyses for RNA-seq: transcript-level estimates improve gene-level inferences. F1000Research. 2015;4:1521.

76. Lawrence M, Gentleman R, Carey V. rtracklayer: an R package for interfacing with genome browsers. Bioinformatics. 2009;25:1841–2.

77. Love M, Anders S, Huber W. Differential analysis of count data–the DESeq2 package. Genome Biol. 2014.

78. Huang DW, Sherman BT, Lempicki RA. Systematic and integrative analysis of large gene lists using DAVID bioinformatics resources. Nat Protoc. 2008;4:44–57.

79. Kolde R. pheatmap: Pretty Heatmaps. 2018.

80. Wickham H. ggplot2. Wiley Interdiscip Rev Comput Stat. 2011;3:180–5.

81. Khamis AM, Hamilton AR, Medvedeva YA, Alam T, Alam I, Essack M, et al. Insights into the Transcriptional Architecture of Behavioral Plasticity in the Honey Bee Apis mellifera. 2015;5:11136.

