## Supplementary figures and images for "Honey bee (*Apis mellifera*) larval pheromones regulate gene expression related to foraging task specialization"

### Supplemental Figure 1

**Cluster dendrogram with AU/BP values (%)**

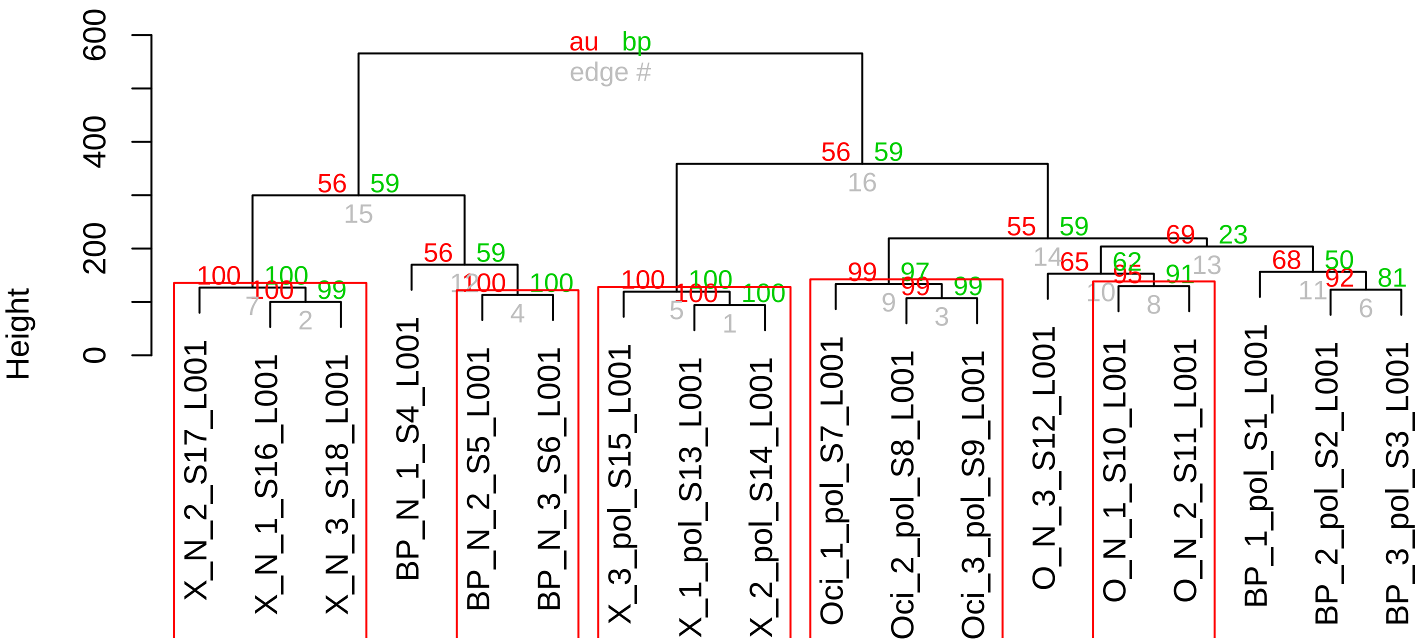


Distance: manhattan
Cluster method: ward.D2

### Supplemental Figure 2

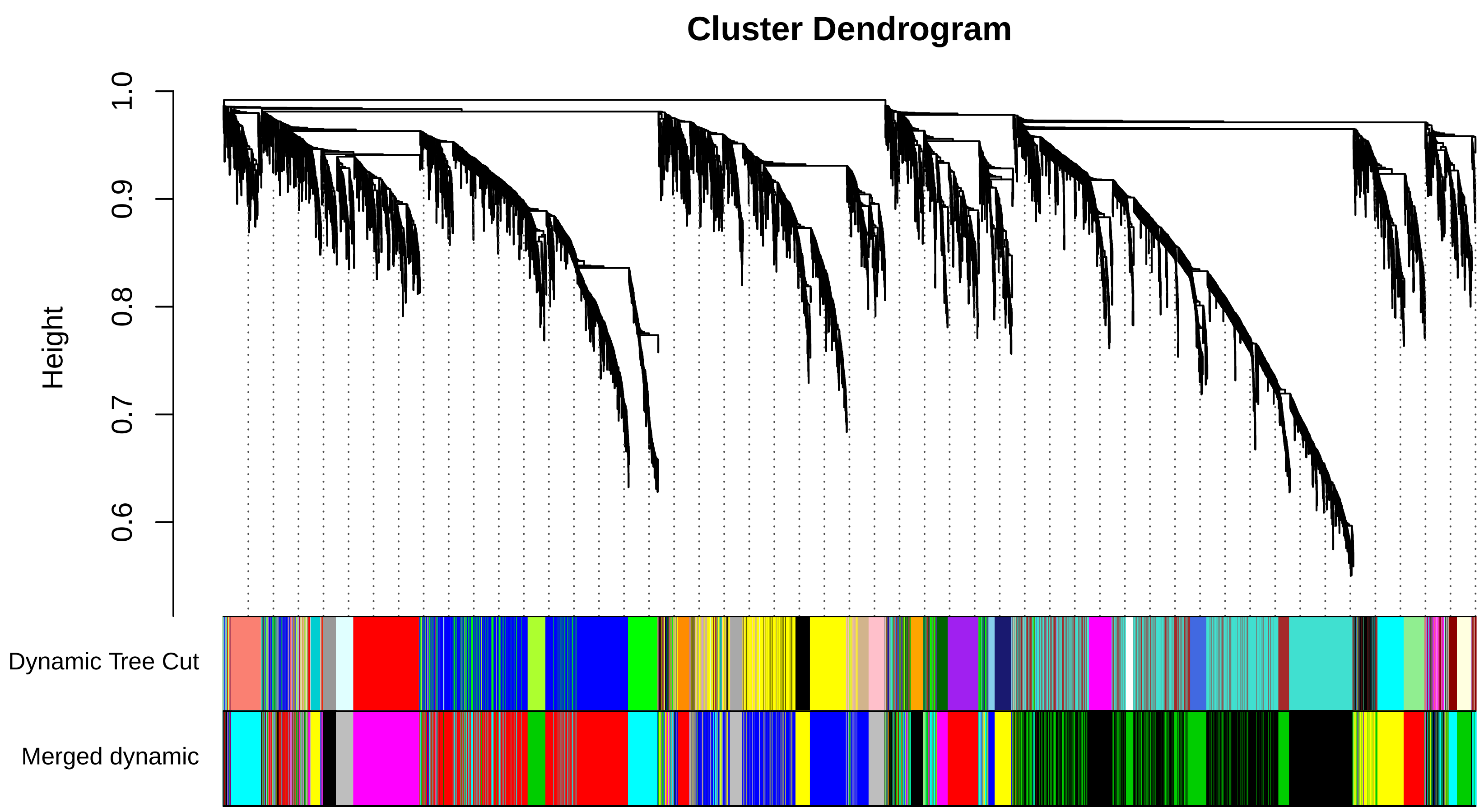
